# Whole Cell Phenotypic Screening Of MMV Pathogen Box identifies Specific Inhibitors of *Plasmodium falciparum* merozoite maturation and egress

**DOI:** 10.1101/772434

**Authors:** Alok Tanala Patra, Tejashri Bhimashankar Hingmire, Meenakshi Belekar, Aoli Xiong, Gowtham Subramanian, Zbynek Bozdech, Peter Preiser, Dhanasekaran Shanmugam, Rajesh Chandramohanadas

## Abstract

We report a systematic, cellular phenotype-based antimalarial screening of the MMV Pathogen Box collection, which facilitated the identification of specific blockers of late stage intraerythrocytic *Plasmodium falciparum* maturation. First, from standard growth inhibition asays, we discovered 62 additional antimalarials (*EC*_50_ ≤ 10μM) over previously known antimalarial candidates from Pathogen Box. A total of 90 potent molecules (*EC*_50_ ≤ 1μM) were selected for evaluating their stage-specific effects during the intra-erythrocytic development of *P. falciparum*. None of these molecules had significant effect on ring-trophozoite transition, 10 molecules inhibited trophozoite-schizont transition, and 21 molecules inhibited schizont-ring transition at 1μM. These compounds were further validated in secondary assays by flow cytometry and microscopic imaging of treated cells to prioritize 12 molecules as potent and selective blockers of schizont-ring transition. Seven of these were found to strongly inhibit calcium ionophore induced egress of *Toxoplasma gondii*, a related apicomplexan parasite, suggesting that the inhibitors may be acting *via* similar mechanism in the two parasites, which can be further exploited for target identification studies. Two of these molecules, with previously unknown mechanism of action, MMV020670 and MMV026356, were found to induce fragmentation of DNA in developing merozoites. Further mechanistic studies would facilitate therapeutic exploitation of these molecules as broadly active inhibitors targeting development and egress of apicomplexan parasites relevant to human health.

## Introduction

Despite technological advancements and improved knowledge on disease manifestation and pathogen biology, progress in therapeutic development has been slower than expected for human infectious diseases affecting the developing world. In addition to limited financial prospects, pharmaceutical companies remain under significant pressure to address high attrition rates. In this context, repurposing of old drugs is becoming a viable alternative through the exploitation of known compounds with established mechanisms thereby shortening the time window between efficacy testing to pharmacophore optimization^1–3^ to eventual application. In recent efforts to speed up the process of drug development against various pathogenic diseases, a collaborative model of public-private partnerships (PPPs) facilitated through Medicines for Malaria Venture (MMV: mmv.org) was formed. This resulted in well-characterized drug-like small molecule libraries being made accessible to academics worldwide, for the purpose of identifying novel bioactivities against various tropical disease pathogens and conducting focused mechanistic studies. In 2015, the consortium launched the open access “Pathogen Box” consisting of 400 drug-like small molecules, included 26 reference compounds, with confirmed bioactivity against at least one of the neglected disease pathogens (*Plasmodium*, *Mycobacterium*, Kinetoplastids, *Schistosoma*, *Cryptosporidium*, helminths and dengue virus),^4^ as a way to facilitate open-ended drug discovery.

Open access to the Pathogen Box has led to the discovery of newly identified activities against various pathogens, for example, MMV688934 (Tolfenpyrad) showed activity against the helminth parasite and barber’s pole worm^5^ while MMV688768 was shown to demonstrate inhibitory activity against *Candida albicans* biofilm formation^6^. Previously, these molecules were established as inhibitors of kinetoplastid and schistosomiasis^5–6^ respectively. However, the latter molecule has been recently documented to have no activity against *S. mansoni*^7^. Dual activity against *Giardia* and *Cryptosporidium* were reported for MMV010576, MMV028694 and MMV676501 that showed ≥ 75% inhibitory efficacy^8^. Several such studies have accelerated the identification of small molecules that are now re-profiled, repurposed and reclassified to show broader inhibitory activities on more than one pathogenic agent^9–12^.

*Plasmodium falciparum* is the most virulent human malaria species and is widespread, accounting for 99.7% of estimated malaria cases in the African Region, 62.8% in South-East Asia, 69% in Eastern Mediterranean region and 71.9% in Western Pacific region^13–15^. Within the human host, *Plasmodium* undergoes vastly different developmental changes in liver cells and red blood cells. However, intra-erythrocytic (blood stage) development is most obvious during infection and is responsible for malaria-associated pathological outcomes. Over the past several decades, many attempts have been made to eradicate malaria but with limited success due to challenges in the form of drug resistance, changes in vector distribution and density, and improper regulatory framework in many malaria endemic areas. Malaria-endemic regions are rapidly losing affordable and effective therapeutic options, making it necessary to develop new drugs with novel modes of action and molecular targets.

We have exploited the MMV Pathogen Box to dissect the blood stage specific antimalarial activity of the molecules in this library. To document the stage-specific activity of MMV Pathogen Box molecules against *P. falciparum*, a phenotype-based screening approach was used. First, growth inhibition assays were carried out to assess the antimalarial potential of all 400 molecules in the library. This revealed the previously known antimalarial compounds as well as 62 additional novel antimalarial compounds from MMV Pathogen Box. Further evaluations on 90 most potent molecules (*EC*_50_ ≤ 1μM) revealed their stage specific inhibitory effect during the course of blood stage *P. falciparum* development. This identified 10 molecules that inhibited trophozoite development and 21 molecules that inhibited schizont-ring transition. Based on drug-induced phenotypes, 12 of these molecules were found to selectively impair maturation and egress of merozoites. Interesting, 7 of these molecules were potent inhibitors of calcium ionophore induced egress of tachyzoite stage *Toxoplasma gondii*, suggesting a likely conserved mechanism of action across species. We demonstrate that two of these molecules affect DNA integrity in developing *Plasmodium* merozoites while the others appear to interfere directly with the egress process. These findings provide starting points for subsequent mechanistic studies on these molecules and further development of new therapies to treat malaria and toxoplasmosis.

## Materials and Methods

### MMV Pathogen Box Chemical Library: Stock solutions and Storage

The MMV Pathogen Box was received in 96-well plates containing 10µl of 10mM stock solution in 100% dimethyl sulfoxide (DMSO). The stocks were immediately diluted and aliquoted to 5 subsets with a final drug concentration of 1mM in DMSO (20µl of stock solution and 180µl of filtered DMSO), as recommended by MMV. These plates were stored at −80° C. Care was taken to avoid multiple freeze-thaw of the plates during the course of this study.

### *P. falciparum in vitro* culturing and maintenance

Blood for culturing *P. falciparum* was purchased from Interstate Blood bank (USA). All experiments reported in this study used the 3D7 strain of *P. falciparum* and infected RBCs were maintained in sodium bicarbonate/Hepes-buffered RPMI-1640 culture medium (Sigma Aldrich, Singapore) at pH 7.4, supplemented with hypoxanthine and 2.5 mg/mL Albumax II (Gibco) [^12^]. Ring stage parasites were synchronized by treatment with 5% sorbitol at 37° C. Parasites used for the schizont-ring transition assays were enriched using MACs separation (Miltenyi Biotech, Bergisch Gladbach, Germany), as reported in prior work^16^. Parasitemia was scored using flow cytometry (Accuri C6 sampler, BD biosciences Singapore) after staining with Hoechst dye (Hoechst 33342, Thermo Fisher). In parallel, Giemsa-stained thin smears were microscopically examined to match with flow cytometry results and to record phenotypic characteristics arising from drug treatment, as established previously^17–18^.

### EC_50_ determination of Pathogen Box molecules against *P. falciparum*

Antimalarial efficacy screening was performed in 96 well plate format. The molecules were plated by 2-fold serial dilution, from 10 µM to 0.01 µM, and the parasite culture (200 µl per well) was maintained at 2.5 % hematocrit and ∼ 2 % starting parasitemia. As positive controls, the antimalarial drugs atovaquone and/or chloroquine were used at 1 µM each, while 1 % DMSO treatment served as the negative control. Parasite growth inhibition was estimated after 60 h incubation of assay plates under optimal growth condition. For assay readout, 25 µl of staining reagent (10X SYBR Green dye and 0.5% Triton X-100 in phosphate buffered saline (PBS), pH 7.4) was mixed into each well of assay plate and fluorescence emitted by DNA bound dye was quantified, using excitation and emission wavelengths of 498 nm and 522 nm respectively, in a GloMax plate reader (Promega). Data processing and statistical analysis were performed using Microsoft Excel software.

### *In vitro* assessment of stage-specific inhibition

A total of 90 MMV Pathogen Box molecules with nanomolar potency (*EC*_50_ ≤ 1μM), were chosen for testing their inhibitory effect on the development and progression of ring stage (6-8 hours post invasion (hpi)), trophozoite stage (22-24 hpi) and schizont stage (40-42 hpi) *P. falciparum*. The parasites were treated with the inhibitors for 12-14 h in each stage, following which the cultures were stained with Hoechst 33342 and analyzed by flow cytometry to estimate parasitemia and determine the parasite stage^17^. Briefly, 50 µl of culture was aliquoted from the master assay plates and fixed with 0.1 % glutaraldehyde in PBS at 4° C overnight. The cells were then washed once with PBS and permeabilized with 0.25 % Triton X-100 for 10 min at room temperature and washed twice with PBS before staining with Hoechst 33342. Flow cytometry was performed using the Accuri C6 instrument (BD Biosciences), and 100,000 events was recorded and analyzed for deriving parasitemia and parasite stage transition data. Statistical analysis of the data was performed using GraphPad Prism (GraphPad Software, Inc.)^19^.

### Characterization of drug-induced phenotypes and rupture50 (R_50_) determination

Following inhibitor exposure, thin blood smears were prepared on glass sides which were then fixed using 100% methanol and stained with 1:10 ratio filtered Giemsa (Merck Millipore) in distilled water. A standard phase-contract light microscope (Leica DM750, Leica Microsystem) was utilized for visualising the phenotypic changes of the cell and image capturing. For R_50_ determination, synchronized schizont stage parasites (∼40 to 42 hpi) were treated with selected inhibitors at 10, 3, 1, 0.3 and 0.1 µM concentration for 12 h, at which point ring stage parasites can be observed in DMSO-treated control cultures. Blood smears were taken for staining with Giemsa at various time points. In parallel, cells were harvested for flow cytometry analysis for further validation. DMSO and E-64 (a broad spectrum cysteine protease inhibitor known to completely block *P. falciparum* egress were used as controls for all egress studies^20^.

### Measurement of Intracellular ROS

Magnet purified schizont stage parasites (∼42-44 hpi) were treated with selected Pathogen Box molecules at a concentration of 1 µM for 1 h. Following incubation, 5 μM CellROX® Green reagent (Thermo Fisher Scientific, Inc.) was added to the cells and incubated for 30 min in a 5% CO_2_ incubator at 37° C. The cells were co-stained with Hoechst 33342 during the last 15 min of incubation in MCM at 37° C in dark. The cells were then washed with PBS three times and visualized using a fluorescent microscope (Olympus CKX53 inverted microscope).

### Assessment of DNA damage using Comet Chip

Alkaline Comet Chip assay was performed as described ^21–22^. Briefly, schizont stage *P. falciparum* parasites were enriched on a percoll gradient and resuspended in complete RPMI at 0.5% v/v. Cell suspensions were incubated with different concentration of drugs or mock solution (0.05% DMSO) in 96-well plate for 1 h at 37 °C. Treated cells were transferred to Comet Chip, lysed overnight at 4°C in alkaline lysis buffer and electrophoresis at 4°C for 30 min at 1 V/cm and ∼300 mA with alkaline electrophoresis buffer as previously described ^21^. The chip was then washed twice at room temperature with neutralization buffer containing 0.4M Trizma® HCl (Sigma-Aldrich, USA). The Comet Chip was stained with 1X of SYBR™ Gold (Invitrogen, USA) for 30 min at room temperature in dark. Fluorescent images of the comets were captured at 40X magnification using epifluorescence microscopes, Olympus IX83, with a 480-nm excitation filter. Images were analysed using Guicometanalyzer, a customized software developed in MATLAB (The MathWorks Inc., USA) as previously described ^22^. The median of comets in each well was calculated from results generated in Guicometanalyzer and converted into excel file using Comet to excel, a program developed in Python.

### Egress inhibition studies in *Toxoplasma gondii*

Tachyzoite stage RH strain *T. gondii* were cultured in the laboratory as previously reported^18^. When exposed to the calcium ionophore A23187, intracellular tachyzoites are known to rapidly rupture the parasitophorous vacuole and egress out of the host cells^23–25^. Here, the ionophore induced egress of *T. gondii* tachyzoites was performed after treating the parasites with selected inhibitors (at 10 µM for 24 h) to assess their effect on the egress process. The time required for vacuole rupture and parasite egress was monitored using a 40X objective fitted to an inverted bright-field microscope (Primo Vert, Zeiss) for a period of 10 min and images were captured at 30 sec intervals.

## Results and Discussion

### Antimalarial activity profiling of MMV Pathogen Box Molecules

To first identify all MMV Pathogen Box compounds that possess antimalarial activity, a standard end point growth inhibition assay was performed on blood-stage *P. falciparum* parasites. Sorbitol synchronized ring stage parasites were treated with all 400 Pathogen Box compounds at 10 µM for 60 h and those which inhibited parasite growth by ≥ 80% were selected for *EC*_50_ determination. In the dose assay, the compounds were tested at concentrations ranging from 10 µM to 1 nM. Out of 400 Pathogen Box compounds, 162 molecules (∼40% hit rate) were found to inhibit growth of blood stage *P. falciparum* parasites by ≥ 80%. Out of the 125 Pathogen Box compounds with previously reported antimalarial activity, only 100 showed ≥ 80% inhibition of parasite growth **(Fig. 1A)**, and the remaining 25 compounds had varying levels of inhibitory activity, presumably due to differences in the assay conditions ^10^ or parasitic stages tested. A compilation of screening data is provided in the supplementary information (**Supplemental Dataset, S1)**. Interestingly, 62 compounds, previously shown to be active against other pathogens, were also found to have potent antimalarial activity **(Fig. 1B)**. For instance, anti-kinetoplastid molecules like MMV688362 and MMV688271 were found to inhibit *P. falciparum* growth with nanomolar potency (*EC*_50_ ≤ 0.4 µM). These molecules are predicted to bind to the minor groove at AT-rich regions of DNA in other organisms ^26–27^, which may render them inhibitory to multiple pathogens. Furthermore, MMV652003 and MMV688283, which are benzamide 4-quinolinamine class of compounds and predicted to target leucyl-tRNA synthetase and βeta-hematin formation respectively ^28–29^, were identified positively as antimalarial hits. Another compound, MMV675968, which has chemical features similar to inhibitors of dihydrofolate reductase (DHFR) in *Cryptosporidium and Toxoplasma*^8, 11^, displayed potent antimalarial activity with an estimated *EC*_50_ of 0.07 µM. Therefore, it is likely that this compound may be targeting the DHFR enzyme in *P. falciparum*^30^. MMV671636, a Quinolinone class molecule with anti-filarial activity, showed very potent antimalarial activity with an estimated *EC*_50_ of 0.01 µM. This compound is predicted to target mitochondrial cytochrome bc1 complex, making it a promising candidate for chemoprophylaxis treatment of malaria^31–32^. Compounds previously shown to be active against *Mycobacterium tuberculosis*, such as MMV021660 (guanidine) and MMV687765 (pyrimidine) with *EC*_50_ value of 0.07 and 0.47 µM, were found to be inhibiting *P. falciparum*. Although, their mechanism of action against *P. falciparum* remains unknown, these molecules are known to disrupt the folate pathway and tyrosine kinase activities in *M. tuberculosis*^33–34^. Several other compounds such as MMV688943, MMV688371, MMV688274, MMV688754, MMV688273, MMV688550, MMV023969, MMV495543, MMV637229, MMV668727, MMV676063, MMV688761, MMV688763, MMV688552, MMV688509 belonging to kinetoplastids, tuberculosis, hookworm infections, filariasis, schistosomiasis, toxoplasmosis disease sets in Pathogen Box, showed submicromolar to micromolar efficacy against *P. falciparum* and warrant further consideration.

**Figure 1:**
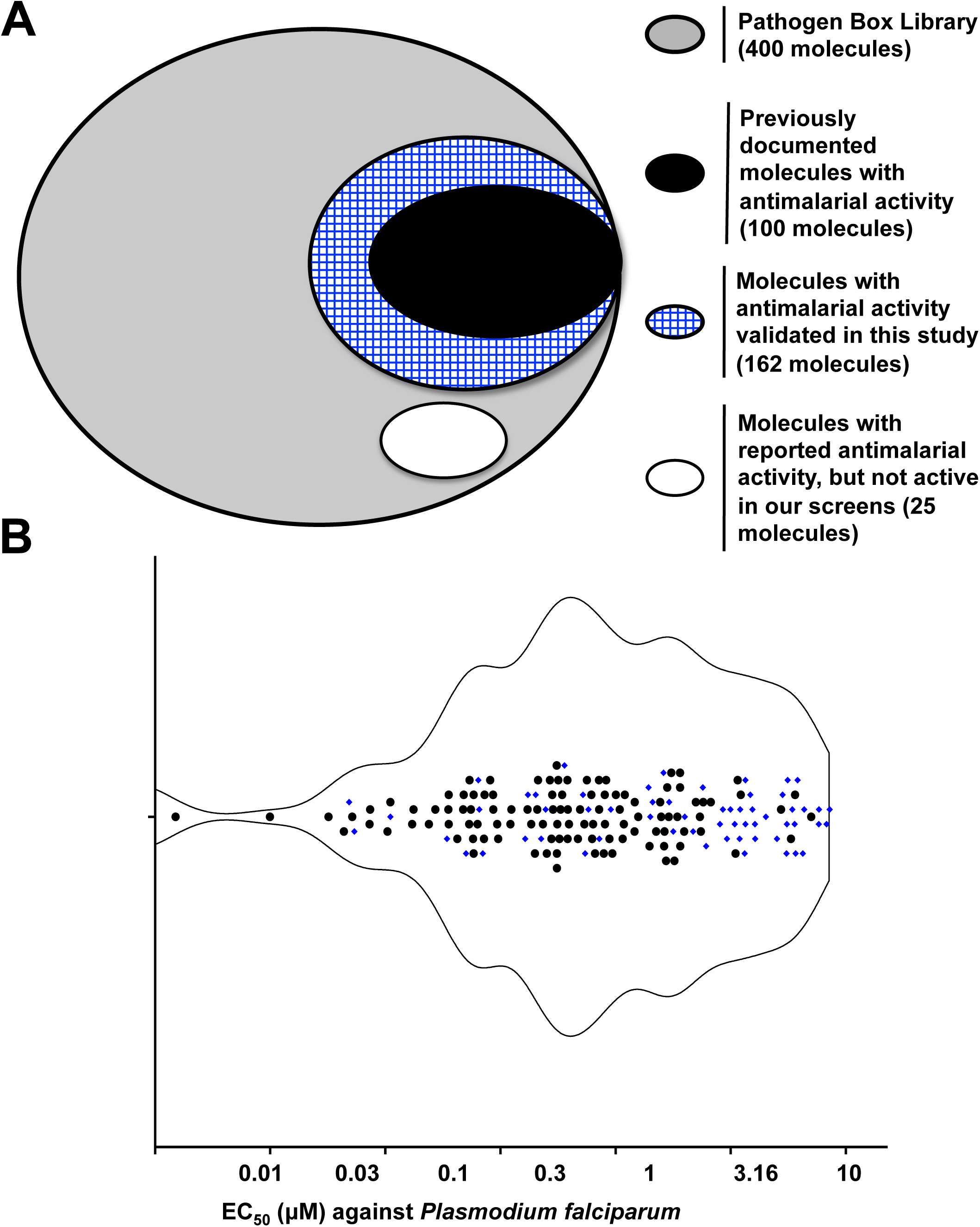
Identifying Pathogen Box molecules inhibiting growth of blood stage *P. falciparum* parasites. (A) Segregation of molecules based on their antimalarial activity. From the 400 molecules tested (indicated in grey), 162 molecules inhibited parasite growth by ≥ 80% at 10 µM concentration (indicated in blue pattern). Out of these, 100 molecules were previously documented for antimalarial activity (indicated in black) and 62 molecules were newly identified in this study. 25 molecules previously reported to have antimalarial activity showed moderate potency (≥50%) in our assays (indicated in white). (B) *EC*_50_ values for the 162 molecules with antimalarial activity are presented to show their antimalarial potency distribution. Newly identified antimalarial molecules from this study are indicated in blue colour.

### Segregation of Pathogen box molecules by stage-specific inhibition of *P. falciparum*

Conventional growth inhibition assays on *P. falciparum* do not capture parasite stage specific effects of small molecule inhibitors. This remains a challenge since drug sensitivity could vary substantially during the 48 h blood stage development lifecycle of *P. falciparum* and can be further influenced by metabolic fate of the drugs under prolonged *in vitro* conditions leading to alterations to the pharmacodynamic properties^35–36^. Therefore, to further categorize the Pathogen Box molecules by stage-specific inhibitory potential, the 90 most potent molecules (*EC*_50_ ≤ 1 µM) were analyzed by phenotype screening. Progressive developmental stages of parasites – rings (6-10 hpi), trophozoites (20-22 hpi) and schizonts (40 to 42 hpi) – were treated with 1 µM inhibitor and assessed after ∼12h treatment for their ability to block stage transition using flow cytometry based detection of the parasite stages **(Fig. 2A)**. None of the molecules showed any significant inhibition of ring-trophozoite transition [R→T], which is in line with previous reports suggesting that ring stage parasites are rather resilient to antimalarial compounds^37–38^. 20 newly identified molecules with EC_50_ value ≤ 1 µM **(Fig. 2A, highlighted in blue)** were part of the stage-transition inhibition assay which showed that these molecules affect different developmental stages of the parasite. We observed varying degree of inhibition for trophozoite-schizont [T→S] and schizont-ring [S→R] transitions for the molecules. For instance, we observed that MMV688271 (guanidine) and MMV671636 (quinolinone) had no effect on [T→S], although they are known to target mitochondria cytochrome bc-1 complex and AT rich sites on the DNA, respectively. Another two molecules, MMV688754 and MMV675968, for which mechanistic insights are limited, were found to block [T→S] and [S→R] transitions. Since trophozoites are engaged in a variety of metabolic activities essential for parasite survival and proliferation, they may offer more target options when compared to the metabolically more quiescent ring stage parasites. Furthermore, trophozoites are also transcriptionally more active and are engaged in preparation of the parasite cell for daughter cell formation and cell division, which makes them more “target-rich” for small molecule inhibitors^39–40^. Interestingly, we identified 21 molecules, which had significant effect on [S→R] transition. Among them, MMV021660 is known to inhibit folate pathway in *Mycobacterium*^41–42^. It is noteworthy that although *EC*_50_ values for some of the newly identified antiplasmodial molecules (MMV637229, MMV024311, MMV687812, MMV687765, MMV652003, MMV676600, MMV688271, MMV688279, MMV688362) are in nanomolar range, their stage specific inhibitory efficiencies are quite varied and distinct. Our observations highlight that for most inhibitors, the overall potency of the inhibitor (expressed as *EC*_50_) as determined from 60 h growth inhibition assays, is poorly correlated to their effect on each stage of malaria parasite. The ability of ring stage parasites to tolerate drug exposure and facilitate emergence of drug resistance is documented^39^. Thus, it is necessary to carefully study the progressive developmental response following inhibitor treatment.

**Figure 2:**
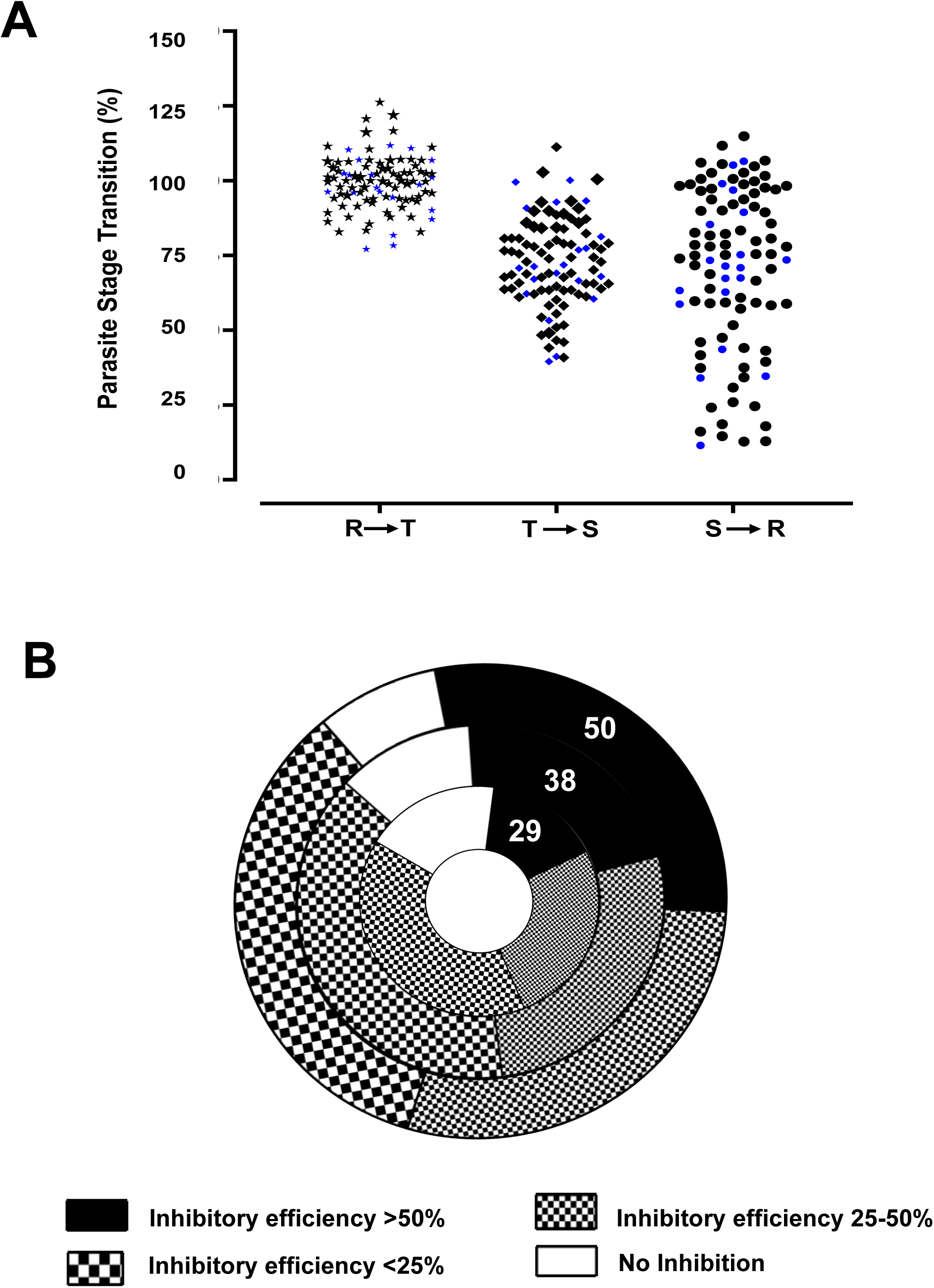
Stage-specific segregation of antimalarial activity for selected pathogen box molecules. **(A)** Flow cytometry data showing % of parasites which have successfully transitioned from ring to trophozoite [RàT], trophozoite to schizont [TàS], and schizont to ring [SàR]. Highlighted in blue are the newly identified molecules with antimalarial activity from this study. **(B)** Pie chart showing inhibition of *P. falciparum* schizont to ring transition upon treatment with 10, 3 and 1 µM of drug concentration (showing inhibitory efficiency <25%, 25 to 50%, >50% and no inhibition). At 10 µM nearly 40% of molecules affected [SàR] transition, which was reduced to 10 % and 5% at 3 and 1 µM respectively, indicating progressive increase in selectivity of the hits identified from higher to lower concentration of the molecules.

### Validation of molecules affecting schizont to ring transition

From the stage transition assays, we observed that a 22 of the screened molecules were inhibiting the [S→R] transition. In order to avoid missing out on other potential [S→R] inhibitors from the antimalarial set of Pathogen Box, we screened all 162 molecules against schizonts. In order to obtain a more detailed insight on the inhibitory activity of the molecules on egress and invasion processes, we employed two levels of validation using flow cytometry and microscopic examination. Synchronized parasites were allowed to develop into schizont stage (40 to 42 hpi) parasites and then treated with three different concentrations (1, 3, and 10 µM) of the inhibitors in the initial screen. After 12 h treatment (i.e., ∼55 hpi) the number of parasites present as newly formed rings and un-ruptured schizonts were scored using flow cytometry. DMSO and E64 were included as negative and positive controls respectively. To observe the parasite cell phenotypic following inhibitor treatment, microscopic examination of Giemsa-stained smears was performed. We observed that 50 of the 162 pathogen box molecules showed inhibitory effect of ≥50% in [S→R] transition at 10 µM, which dropped to 38 molecules at 3 µM and 29 molecules at 1 µM **(Fig. 2B)**. Although 29 molecules inhibited [S→R] transition at 1 µM, microscopic analysis of phenotype indicated that some of them were parasiticidal (and thus unlikely to have any direct effect on the egress/invasion process). Molecules inhibiting [S→R] transition only at 3 µM and 10 µM were not consider for further study since at higher concentrations the observed inhibition can be due to general cytotoxicity on merozoite viability^43–44^.

From our experiments, 12 molecules were short-listed as potent [S→R] transition inhibitors at ≤ 1 µM **(Fig. 3A)**. The following molecules: MMV020520, MMV020710, MMV020391, MMV006239, MMV020623, MMV675958, MMV010576, MMV020670, MMV026356, MMV085071, MMV024443, MMV020081 where efficient inhibitors of schizont maturation at sub micromolar concentrations **(Fig. 3B)**. Microscopy revealed that treatment with the molecules MMV020081 and MMV006239 resulted in cellular phenotypes reminiscent of E64 treatment, indicating specific inhibition of egress process. However, parasites treated with MMV026356 or MMV020670 appeared to be unhealthy and dying, indicating that there may be no direct inhibition of egress or invasion process (**Fig. 3C**). We then performed a concentration-dependent [S→R] transition assay for the top 12 molecules as indicated in **Fig. 3D**. With respect to egress inhibition, MMV020081 appeared to be the most potent with a R_50_ value of 90 nM while its actual estimated *EC*_50_ is 1.7 µM, indicating good selectivity **(Table.1).** Similarly, MMV024443 showed substantially higher potency on egress inhibition with an R_50_ of 0.3 µM in contrast to its *EC*_50_ value of 1.65 µM. MMV675968, which was shown to inhibit thymidylate biosynthesis by targeting the dihydrofolate reductase enzyme in *M. tuberculosis* and *T. gondii*, showed comparable values for *EC*_50_ and R_50_. All of these molecules belong to diverse chemical classes indicating unique mechanism of action (**Supplemental Figure. S1**).

**Figure 3:**
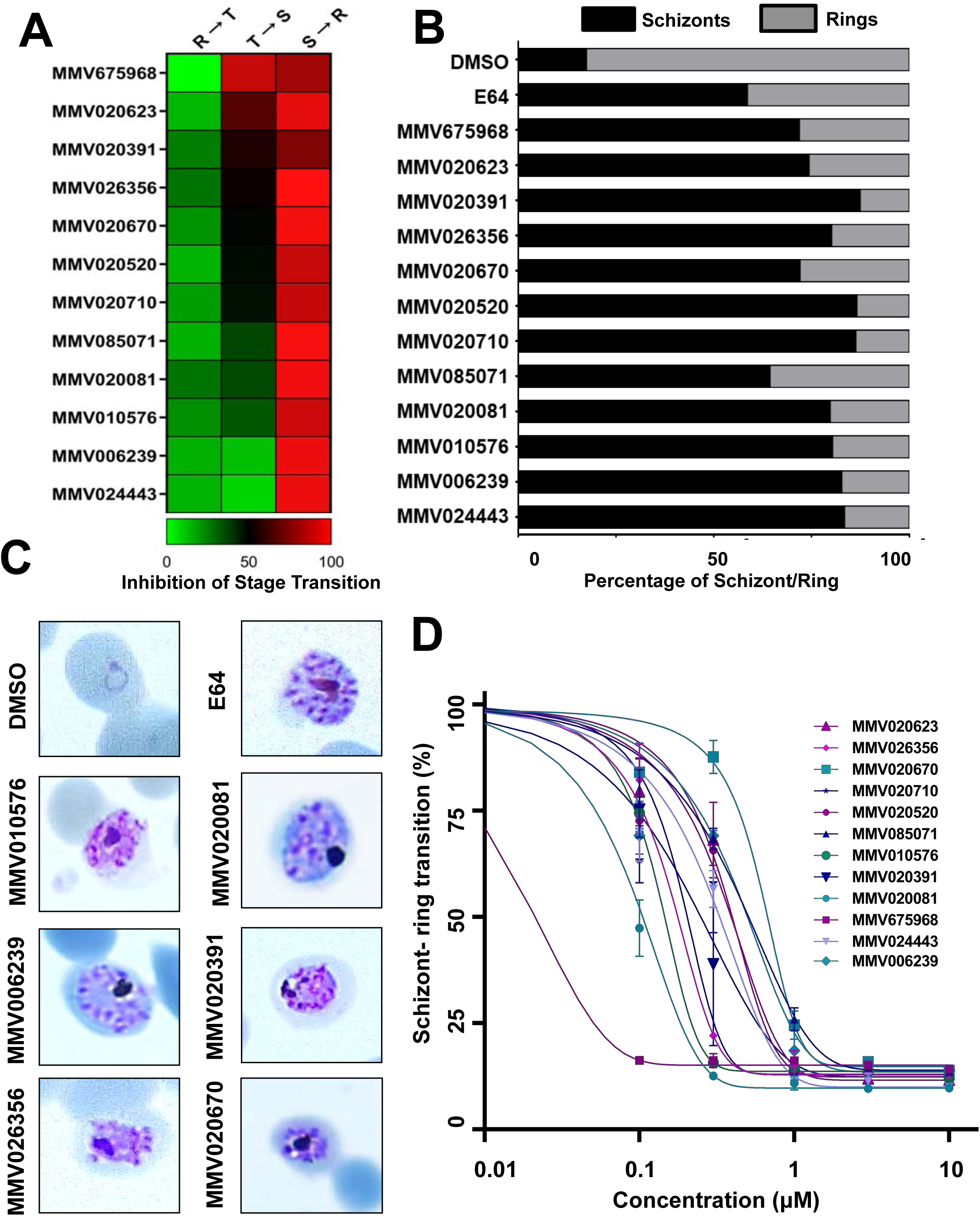
Characterization of schizont inhibitors. **(A)** (Left panel) Showing heatmap for 12 potent schizont inhibitors tested at 1 µM for their stage specific inhibitory effect. While molecules such as MMV675968 and MMV020623 appeared to affect both trophozoite and schizont growth transition, all other molecules were selective against schizont-ring transition **(B)** Late-stage schizonts (42-44 hpi) incubated with the inhibitors were monitored for schizont inhibition through counting newly formed rings by flow cytometry. All molecules tested showed a profound, yet selective activity against [SàR] transition in similar manner to E64 which was included as a positive control for egress inhibition. **(C)** Giemsa-stained smears upon microscopic investigation revealed arrest of late stage parasite development and egress. In case of MMV026356 and MMV020670, unhealthy daughter merozoites were observed indicating cytotoxic activity of these molecules. **(D)** Dose–response curves of potential schizont inhibitors from Pathogen Box library against *P. falciparum* schizonts. DMSO and E64 were included as negative and positive controls respectively. A majority of the Pathogen Box molecules showed remarkable schizonticidal activity.

MMV020623, MMV020520, MMV020081, MMV020710 and MMV020391, which are reported to target PfATP4 and disrupt cellular cation homeostasis, were also found to strongly inhibit [S→R] transition suggesting that PfATP4 activity maybe important for parasite egress^45–46^. MMV024443 (an indole-2-carboxamide) which has been shown to target *P. falciparum* calcium-dependent protein kinase 1 (*Pf*CDPK1)^47–49^ showed excellent egress inhibition in our studies. Previous studies in *P. falciparum* have shown the importance of PfCDPK1 in parasite development and invasion^47–50^. Interestingly, we found that this molecule can also inhibit calcium ionophore induced egress of *T. gondii* (discussed in below section). The *T. gondii* calcium-dependent protein kinase 1 (*Tg*CDPK1) is an essential regulator of calcium-dependent parasite egress and invasion^49^. *Pf*CDPK1 and *Tg*CDPK1 are orthologous and share a sequence identity of 53.85%, suggesting that they may have conserved functions in both parasites. We have determined the Rupture 50 (R_50_) values, defined as the concentration of the molecule at which there is a 50% reduction in newly formed rings compared to the DMSO control. We found that all the 12 molecules tested had R_50_ values in sub-micromolar range.

### Effect of *Plasmodium* schizont inhibitors on toxoplasma egress

Being an obligate intracellular parasite, *T. gondii* actively invades and proliferates inside host cells, within the parasitophorous vacuole. At the end of an intra-vacuolar replicative cycle (or prematurely in response to external perturbations), the parasites will have to egress out and again invade a new host cell to survive and proliferate. Thus, egress and invasion are two critical aspects which are essential for continuity of the replicative cycle. We were interested in finding out whether any of the Pathogen Box molecules, which affected the maturation and egress of *P. falciparum* merozoites, might inhibit egress of tachyzoite stage *T. gondii*. For this we used a calcium ionophore mediated egress assay as previously reported^25^. Intracellular tachyzoite stage parasites were treated with 10 µM concentration of inhibitors for 24 h following they were induced to egress using the calcium ionophore (A23187). As control, DMSO (1%) treated cells were used to estimate the normal egress time taken by parasites to egress out. Of the 12 molecules tested, 9 were capable of inhibiting *T. gondii* egress in ionophore induced egress assay. Average egress time for DMSO-treated parasites was ∼1.07 min. For inhibitor-treated parasites, we considered an average egress time delay of at least 4 min to assign them as potential egress inhibitors in *T. gondii* (**Fig. 4A**). Interestingly, we identified 7 molecules (MMV020081, MMV085071, MMV024443, MMV026356, MMV020670, MMV006239 and MMV020710) with significant egress-blocking activity, and at least in case of 3 molecules (MMV020670, MMV020081 and MMV026356) egress block extended beyond 7 minutes **(Fig 4A, highlighted in blue)**. Only two of the molecules tested, MMV020623 (average egress time, ∼4.06 min) and MMV020520 (average egress time, ∼3.26 min) showed modest inhibition of *T. gondii* egress.

**Figure 4:**
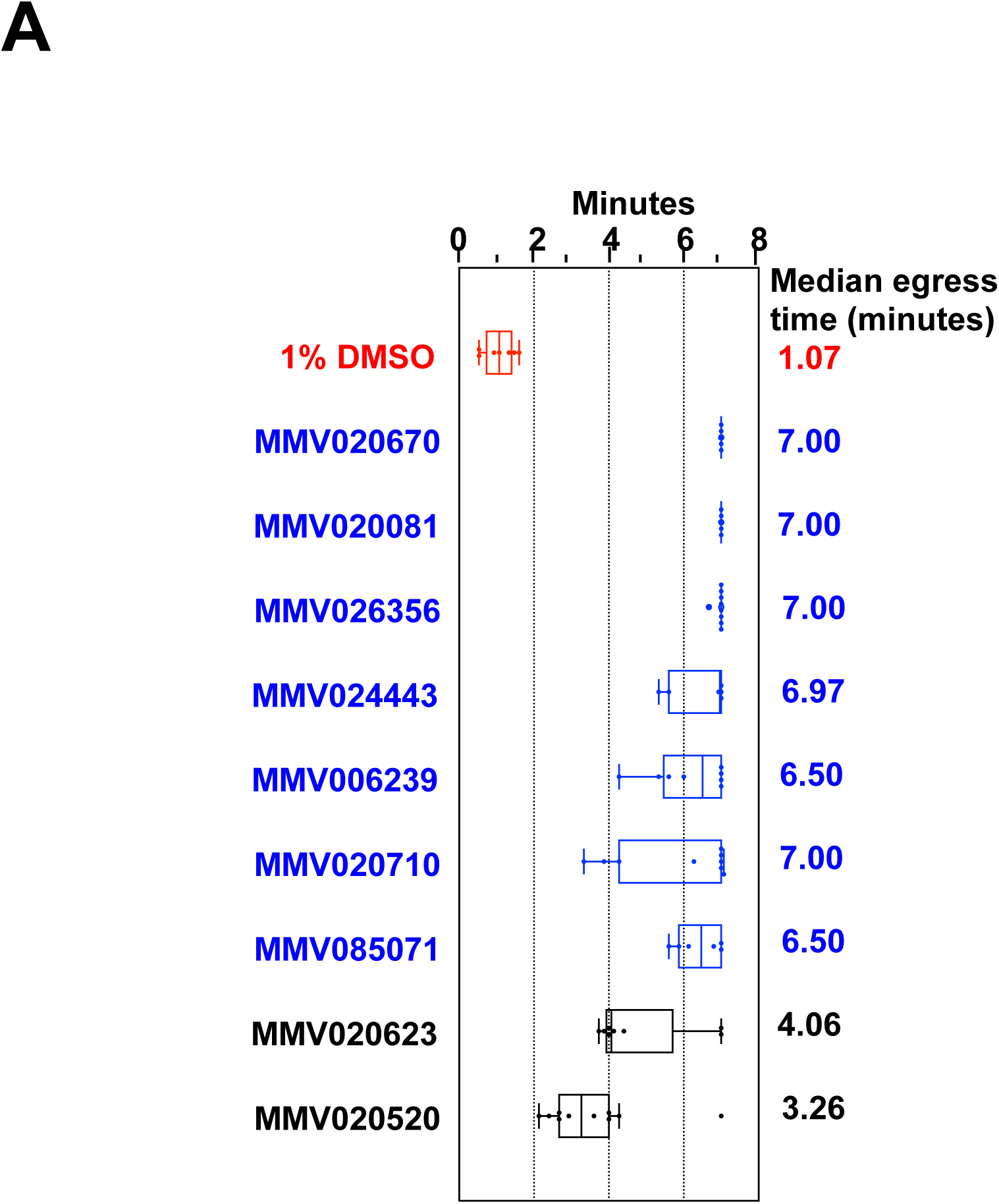
*P. falciparum* [Sà R] blockers inhibit calcium ionophore-mediated egress of *T. gondii* tachyzoites. (A) Whisker plots showing the timing of ionophore-induced egress following 24 h of treatment for Pathogen Box molecules at 10 µM. Red, 1% DMSO-treated control cells; blue, inhibitors for which the median egress time was >6 min.

### Mechanistic insights into parasite death when treated with MMV020391 and MMV010576

Although we identified several molecules with inhibitory potential against late state parasite development and egress, it is reasonable to expect that these molecules have different targets and operate differently at the molecular level, given their structural diversity. Furthermore, microscopy of cellular phenotypes revealed both genuine egress inhibition (MMV020391 and MMV010576) as well as unhealthy and dying merozoites (MMV026356 and MMV020670). Molecules that are cytotoxic could kill parasites by producing ROS and damaging parasite proteins and DNA^51^. To validate such a possibility, we evaluated ROS generation in late-stage schizonts treated with the 7 molecules that were effective against both *Plasmodium* and *Toxoplasma* (MMV020081, MMV085071, MMV024443, MMV026356, MMV020670, MMV006239 and MMV020710). All 7 molecules were positive for ROS generation, albeit to varying degrees **(Fig. 5A)**. Following this observation, we performed a quantitative flow cytometry-based ROS assay on these molecules, which indicated that at least 5 of these molecules (MMV006239, MMV020670, MMV026356, MMV024443 and MMV085071) were potent ROS generators **(Fig. 5B)**.

**Figure 5:**
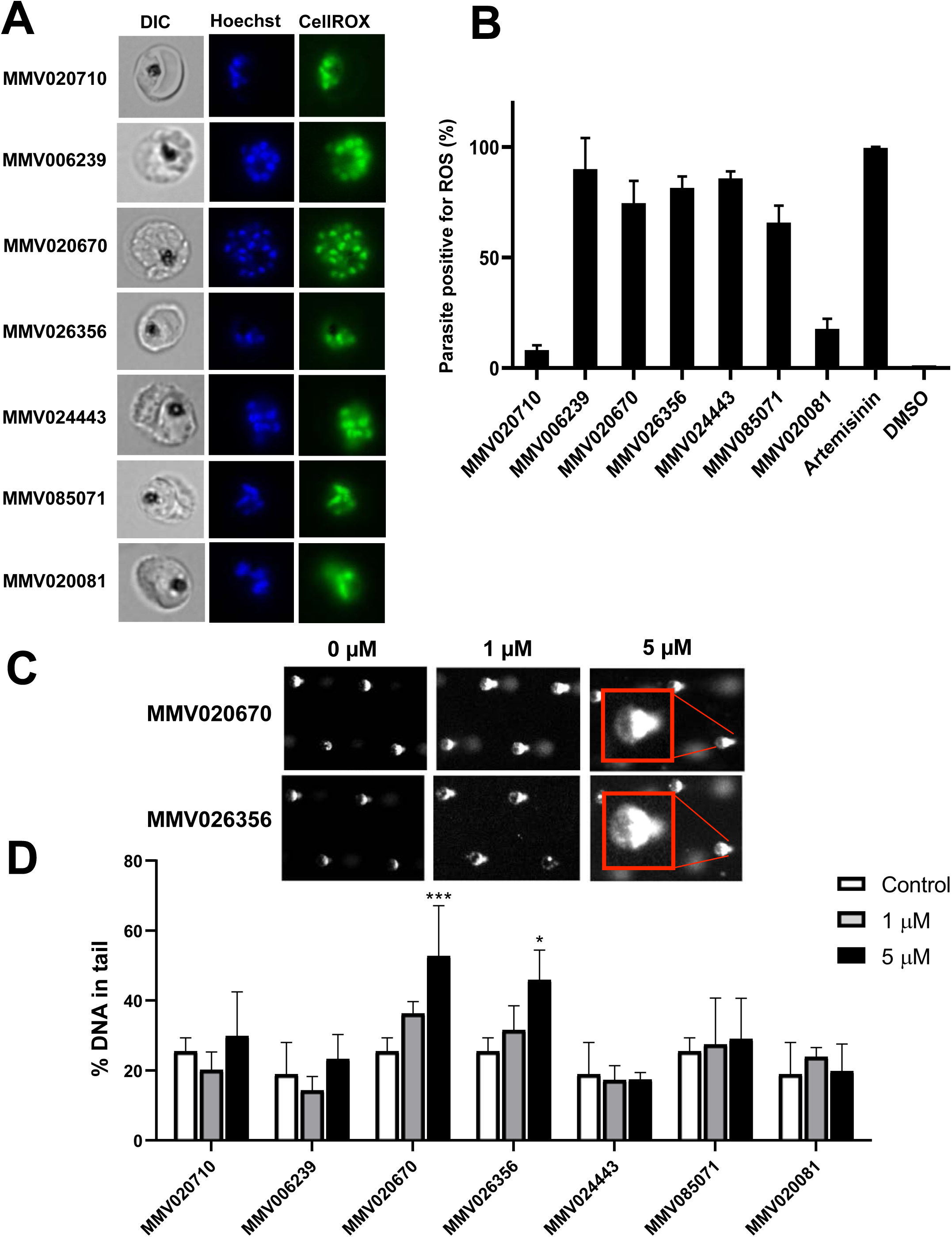
ROS production and DNA fragmentation by Pathogen Box molecules which inhibit parasite egress. (A) Evaluation of ROS production visualized by staining with CellROX Green dye of schizonts after 3 h treatment with 1 µM and 5 µM inhibitor concentration. Varying degrees of ROS production is evident from the observed green fluorescence. (B) Quantification of ROS production using flow cytometry revealed high extent of ROS production for MMV020670, MMV026356 and MMV024443. (C-D) Results from comet assay showing DNA damage in late-stage parasites treated with MMV020670 and MMV026356. None of the other molecules tested showed detectable DNA fragmentation.

This was an interesting observation, and since increased intracellular ROS can cause DNA damage, we tested this using a comet assay that allows visualization of the extent of DNA damage/break-down following inhibitor treatment. From these experiments, ≥50% DNA damage was observed following treatment with the molecules MMV020670 and MMV026356 **(Fig. 5C)**, as indicated by comet tail moment at 5 µM drug exposure **(Fig. 5D)**. Evidently, the increased levels of ROS production generated by MMV020670 and MMV026356 has caused considerable DNA damage. A variety of cellular mechanisms can be responsible for ROS production, including oxidation of lipids, protein and other cellular regulatory mechanisms^52–54^.

Targeting [S→R] transition offers an excellent opportunity for chemotherapy development against *Plasmodium* infection. First, by blocking newly formed invasion ready merozoites from egressing out of infected RBC, rapid increase of the parasite load in clinical cases can be prevented. Second, [S→R] transition involves sequential preparatory and remodelling events in the parasite, which offers numerous targets for inhibitors. Further mechanistic studies on selected Pathogen Box molecules is needed to fully realize the therapeutic potential of these molecules. Studies on altered gene and protein expression, combined with proteome modification studies can be good starting points for obtaining further insights on the mechanism of these molecules.

## Conclusions

Growth inhibition studies on blood stage *P. falciparum* using the MMV Pathogen Box chemical library identified 162 antimalarial molecules, out of which 62 molecules are newly identified. Further, we segregated these molecules based on stage-selectivity, inhibitory efficacy and cellular phenotypes that allowed us to prioritize 12 molecules that interfere with late stage parasite development and egress. We demonstrate that two of the schizont inhibitors, MMV020670 and MMV026356 induce DNA break-down in segmented schizonts leading to parasite arrest. Identifying molecules that are inhibiting egress in both *Plasmodium and Toxoplasma* is very exciting, as this can facilitate target identification and other mechanism studies. This also opens up the possibility of exploiting these molecules a therapeutic agent against multiple apicomplexan parasites.

## Author Contributions

**ATP:** Involved in study design, conducted *Plasmodium* experiments, analysed results, prepared figures and assisted with manuscript preparation, **TH**: Performed antimalarial screening for all Pathogen Box molecules, analysed results and assisted with manuscript preparation, **MB**: Performed *T. gondii* egress assays and analysed results, **AX**: Performed Comet assay analysed results and assisted with manuscript preparation, **GS**: Assisted with plasmodium experiments and assisted with manuscript preparation, **ZB**: Involved in study design, provided tools/reagents and assisted with manuscript preparation, **PRP**: Involved in study design, provided tools/reagents and assisted with manuscript preparation, **DS**: Involved in study design, analysed data, verified results and assisted with manuscript preparation, **RC:** Designed the study, coordinated research, verified results and wrote the manuscript.

## Acknowledgements

We thank MMV for providing the Pathogen Box chemical library used in this research. Authors also than Dr. Kevin S Tan (National University of Singapore) and Dr. DS Reddy (CSIR-National Chemical Laboratories, India) for various discussions.

ATP, GS and RC acknowledge the following grants: RGAST1503 (A*star-India Collaboration Grant) and T1MOE1702 (MoE Tier 1 Grant awarded through SUTD). ATP acknowledges Ministry of Education (MoE), Singapore for President’s Graduate Fellowship. Infrastructure support through SUTD-MIT International Design Centre (IDC) is greatly acknowledged. MAB & TH acknowledge PhD fellowship from the Council of Scientific and Industrial Research, India; DS. acknowledges the Indo-Singapore Joint Science and Technology Research Cooperation grant from the Department of Science and Technology, India (INT/SIN/P-09/2015), and infrastructure support from the CSIR-National Chemical Laboratory, Pune, India. The funders had no role in study design, data collection and interpretation, or the decision to submit the work for publication.

---

**Table 1:**
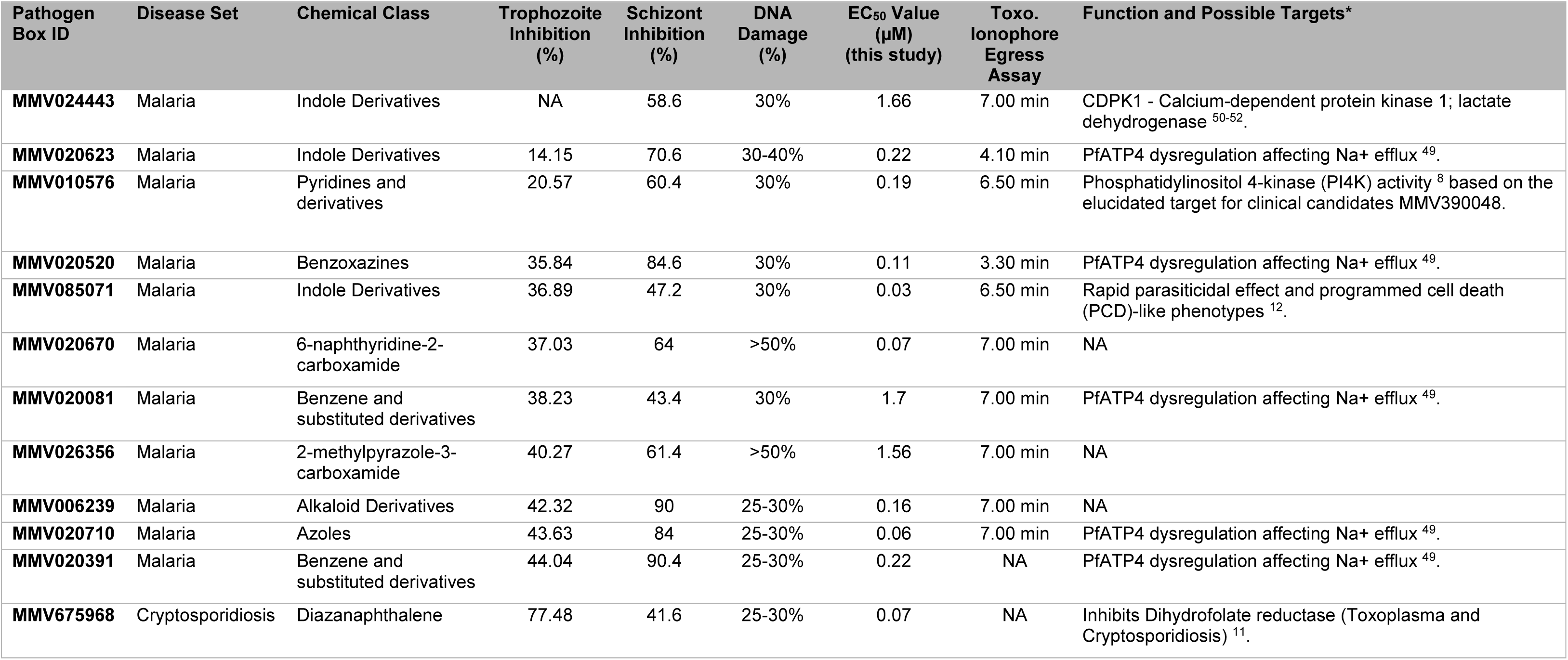
Molecules from other disease sets exhibiting antimalarial activity

| Pathogen Box ID | Disease Set | Chemical Class | Trophozoite Inhibition (%) | Schizont Inhibition (%) | DNA Damage (%) | EC<sub>50</sub> Value (μM) (this study) | Toxo. Ionophore Egress Assay | Function and Possible Targets* |
| --- | --- | --- | --- | --- | --- | --- | --- | --- |
| **MMV024443** | Malaria | Indole Derivatives | NA | 58.6 | 30% | 1.66 | 7.00 min | CDPK1 - Calcium-dependent protein kinase 1; lactate dehydrogenase <sup>50-52</sup>. |
| **MMV020623** | Malaria | Indole Derivatives | 14.15 | 70.6 | 30-40% | 0.22 | 4.10 min | PfATP4 dysregulation affecting Na<sup>+</sup> efflux <sup>49</sup>. |
| **MMV010576** | Malaria | Pyridines and derivatives | 20.57 | 60.4 | 30% | 0.19 | 6.50 min | Phosphatidylinositol 4-kinase (PI4K) activity <sup>8</sup> based on the elucidated target for clinical candidates MMV390048. |
| **MMV020520** | Malaria | Benzoxazines | 35.84 | 84.6 | 30% | 0.11 | 3.30 min | PfATP4 dysregulation affecting Na<sup>+</sup> efflux <sup>49</sup>. |
| **MMV085071** | Malaria | Indole Derivatives | 36.89 | 47.2 | 30% | 0.03 | 6.50 min | Rapid parasitocidal effect and programmed cell death (PCD)-like phenotypes <sup>12</sup>. |
| **MMV020670** | Malaria | 6-naphthyridine-2-carboxamide | 37.03 | 64 | >50% | 0.07 | 7.00 min | NA |
| **MMV020081** | Malaria | Benzene and substituted derivatives | 38.23 | 43.4 | 30% | 1.7 | 7.00 min | PfATP4 dysregulation affecting Na<sup>+</sup> efflux <sup>49</sup>. |
| **MMV026356** | Malaria | 2-methylpyrazole-3-carboxamide | 40.27 | 61.4 | >50% | 1.56 | 7.00 min | NA |
| **MMV006239** | Malaria | Alkaloid Derivatives | 42.32 | 90 | 25-30% | 0.16 | 7.00 min | NA |
| **MMV020710** | Malaria | Azoles | 43.63 | 84 | 25-30% | 0.06 | 7.00 min | PfATP4 dysregulation affecting Na<sup>+</sup> efflux <sup>49</sup>. |
| **MMV020391** | Malaria | Benzene and substituted derivatives | 44.04 | 90.4 | 25-30% | 0.22 | NA | PfATP4 dysregulation affecting Na<sup>+</sup> efflux <sup>49</sup>. |
| **MMV675968** | Cryptosporidiosis | Diazanaphthalene | 77.48 | 41.6 | 25-30% | 0.07 | NA | Inhibits Dihydrofolate reductase (Toxoplasma and Cryptosporidiosis) <sup>11</sup>. |

**Fig S1:**
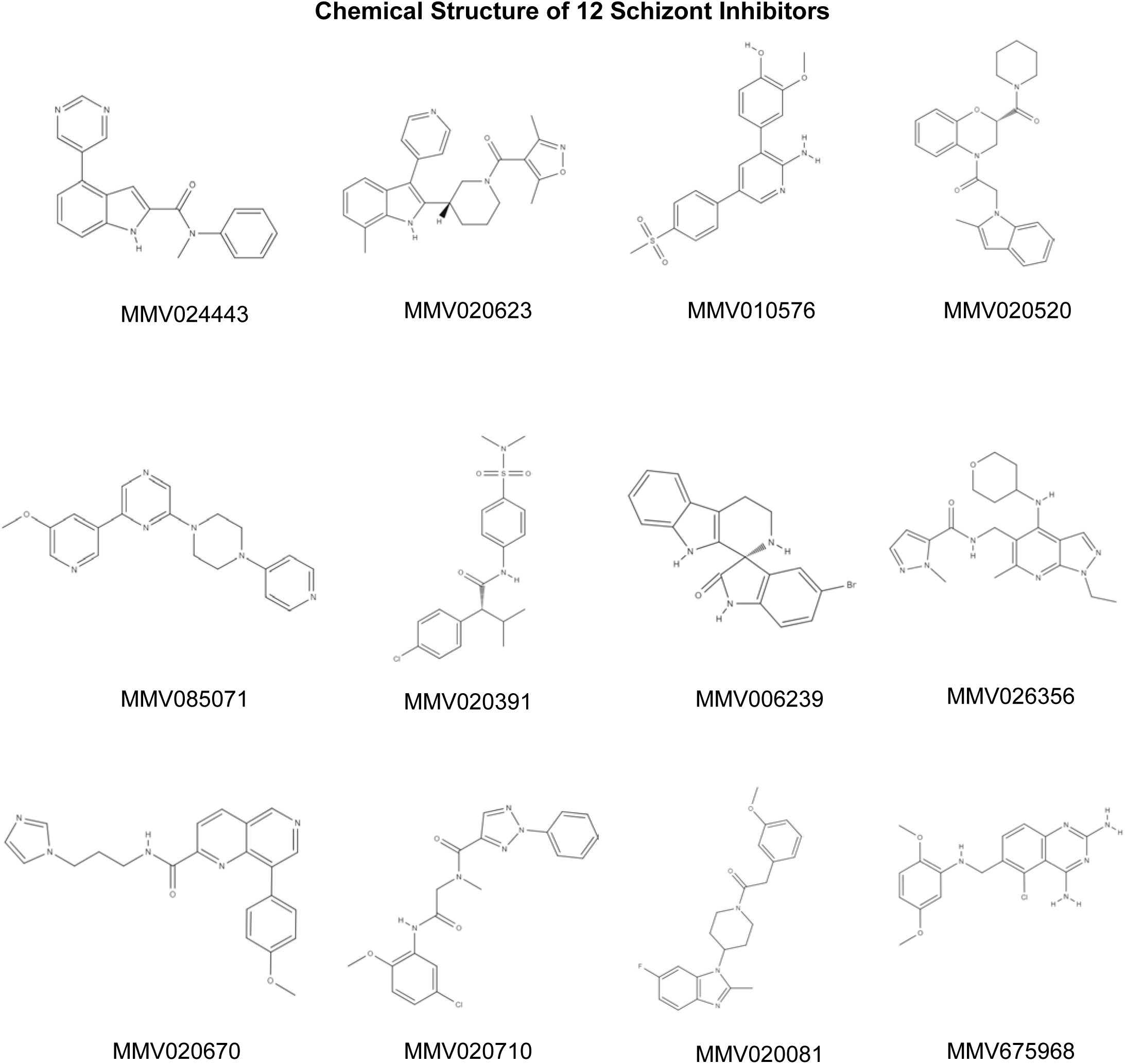
Structure of the 12 most effective compounds identified by *in vitro* screening using 1µM as the highest concentration, as late stage parasite inhibitors. These compound structures are obtained from PubChem (https://pubchem.ncbi.nlm.nih.gov) and MolView (http://molview.org/)

**Fig S2:**
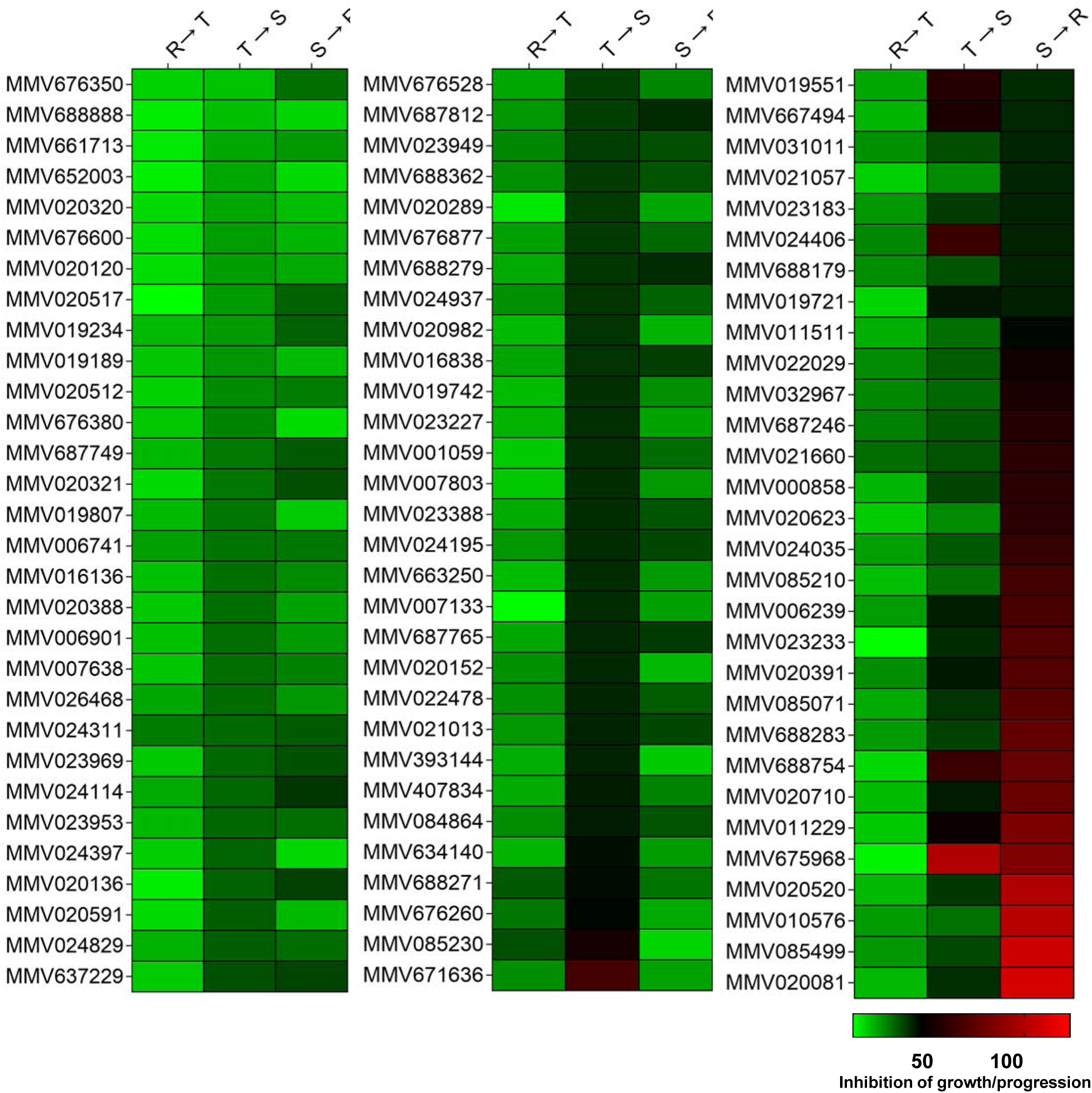
Heat Map showing parasite growth progression data of 96 molecules testing against *P. falciparum*. The data shown are mean values of two biological replicates.

**Table 2:**
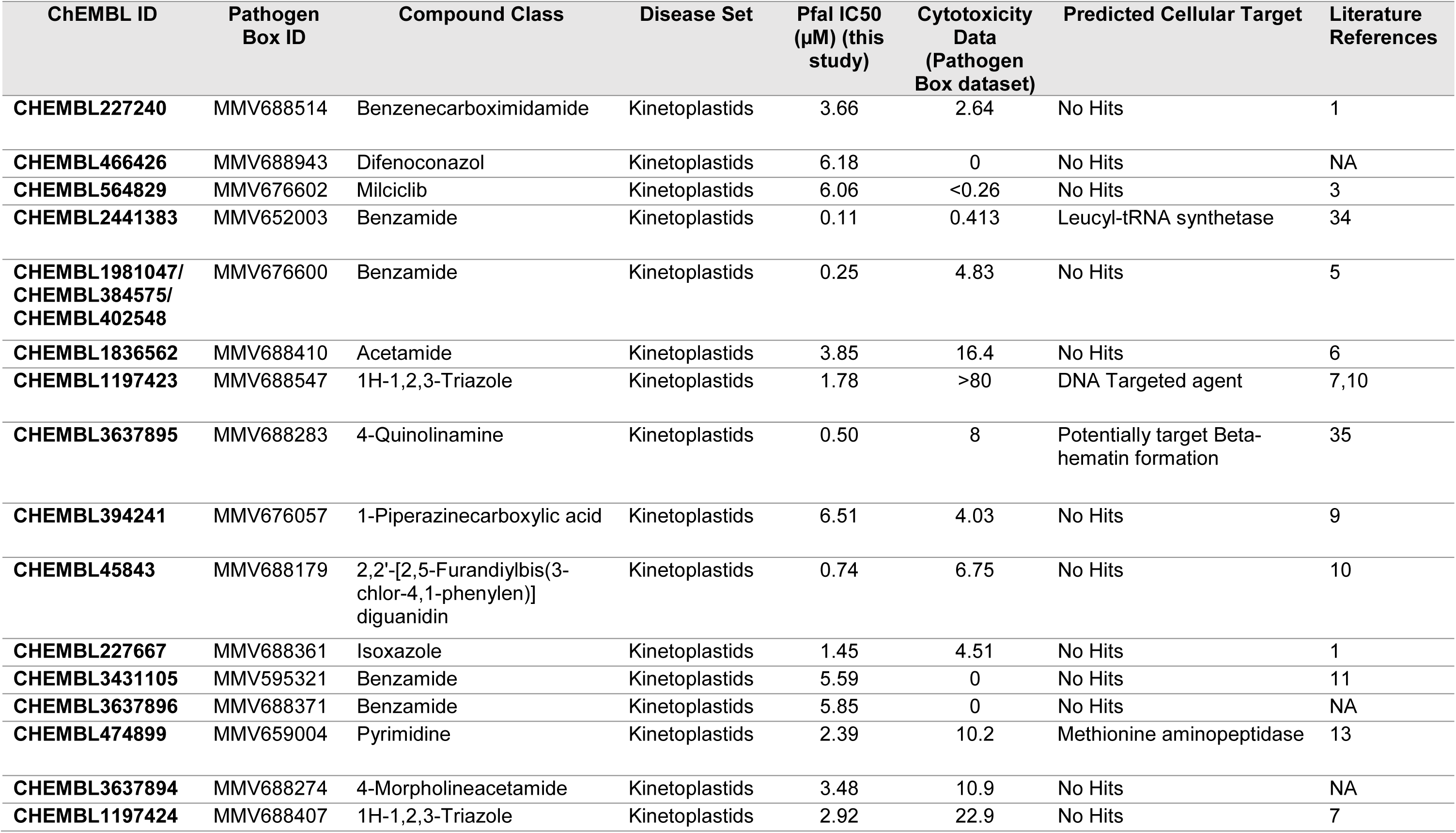
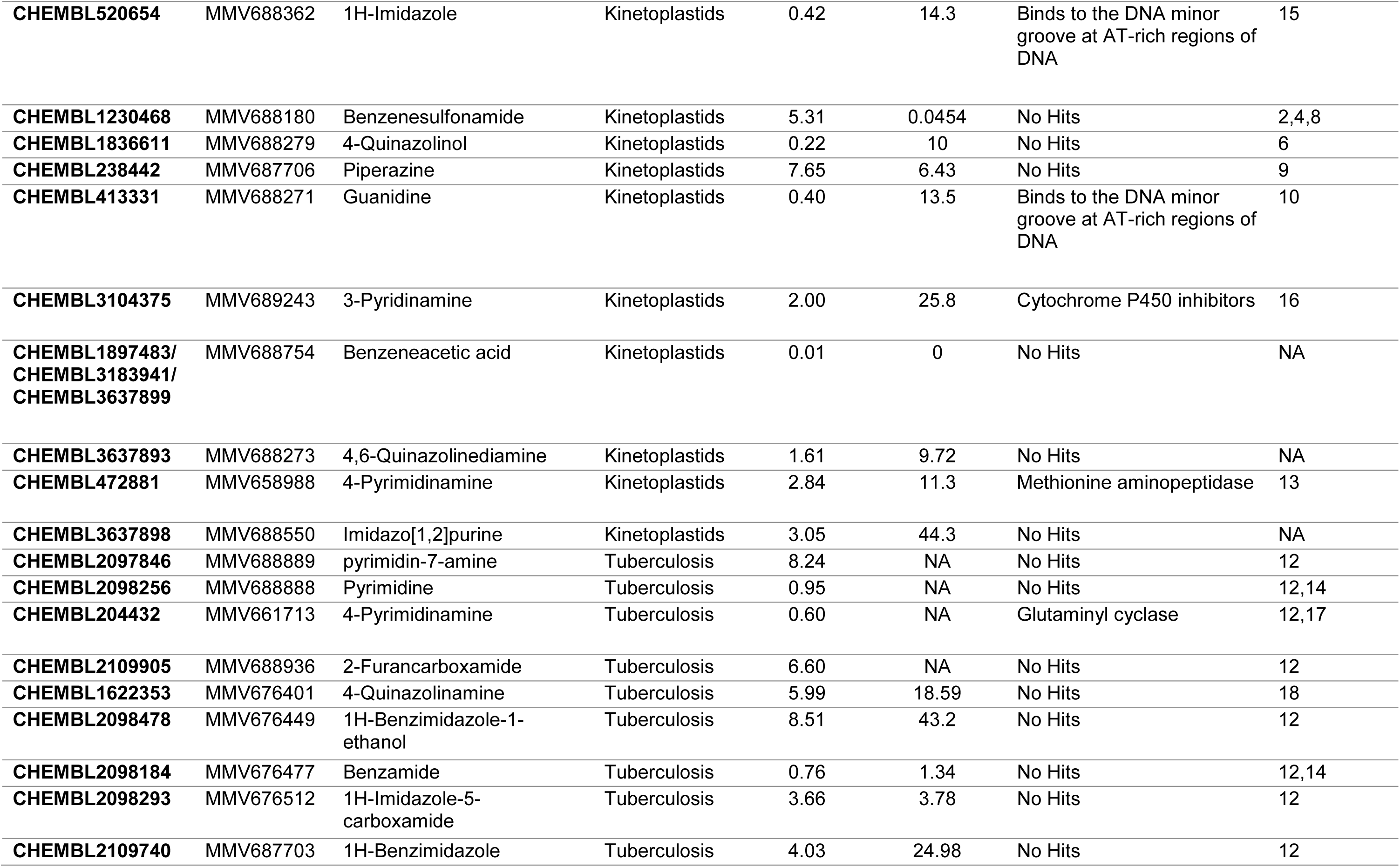
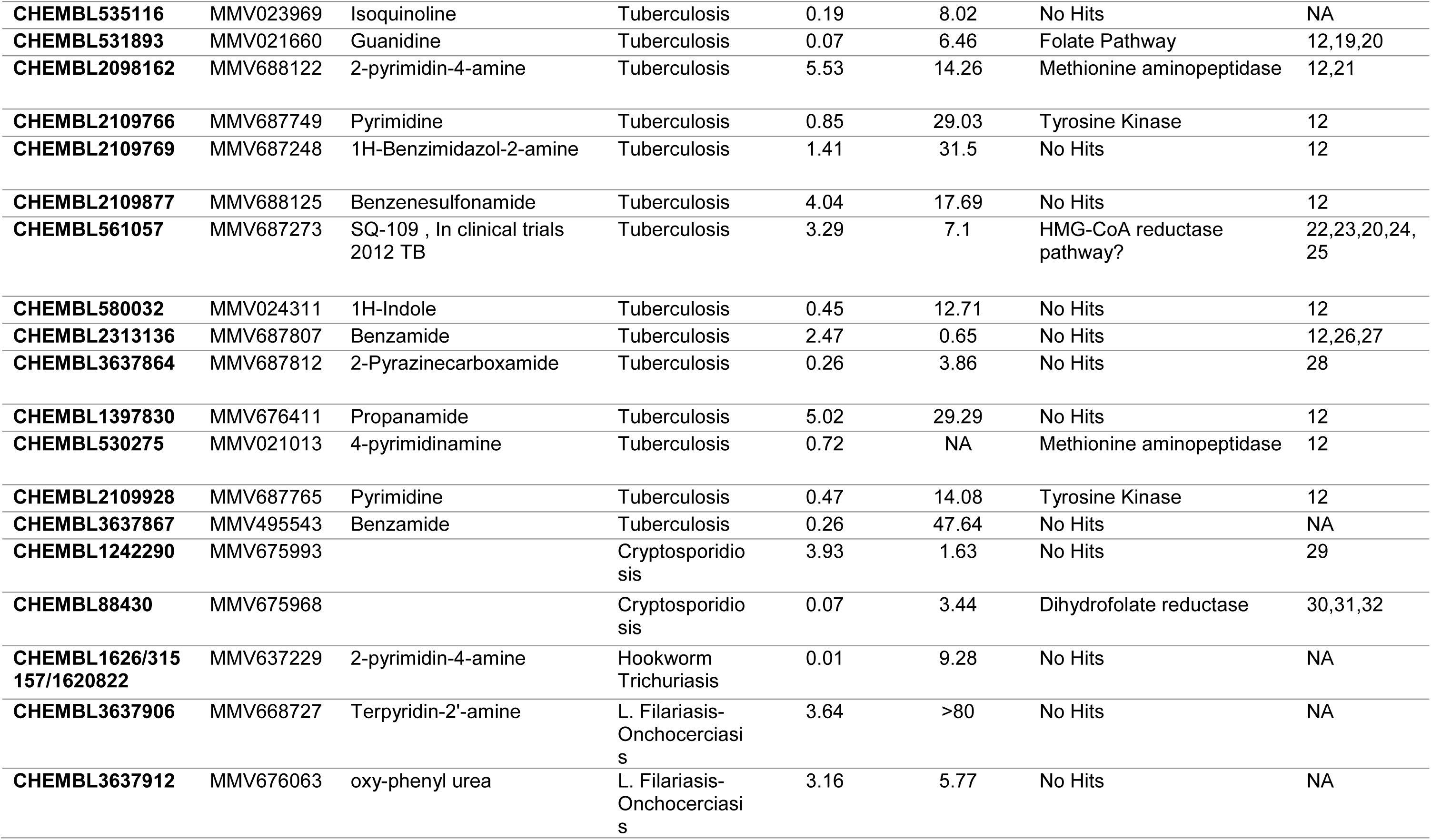
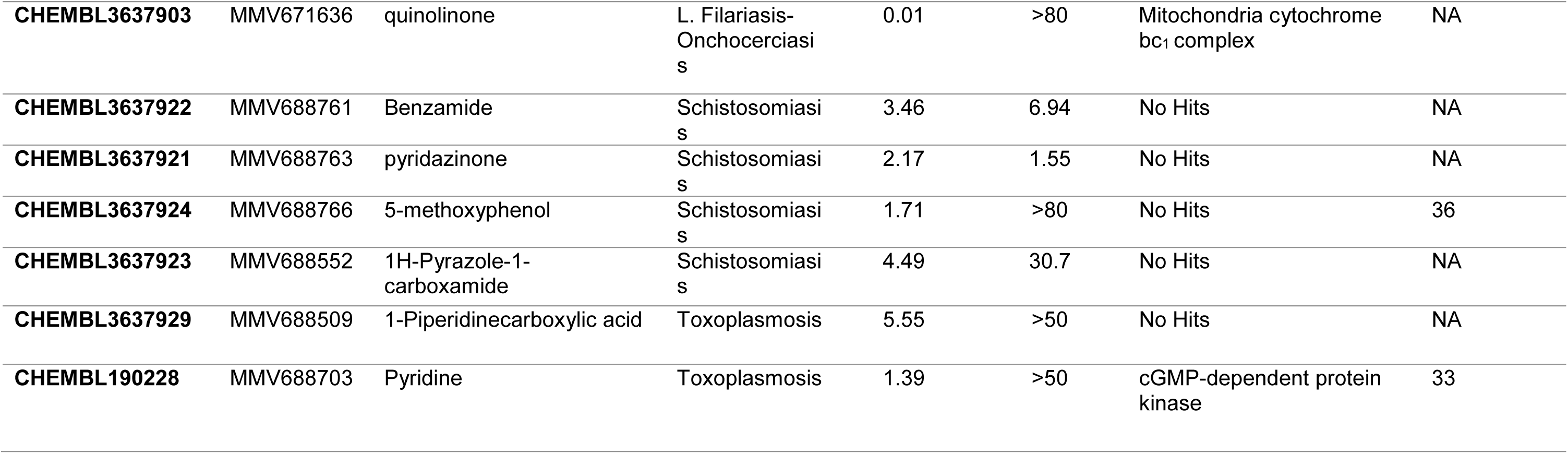
Molecules from other disease sets exhibiting antimalarial activity

