## Supplementary Figure S1 for "Whole Cell Phenotypic Screening Of MMV Pathogen Box identifies Specific Inhibitors of *Plasmodium falciparum* merozoite maturation and egress"

### Fig S1

**Chemical Structure of 12 Schizont Inhibitors**

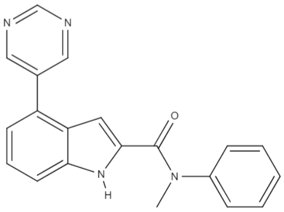

MMV024443

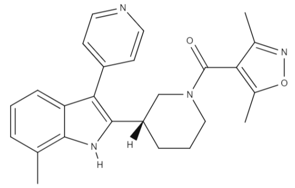

MMV020623

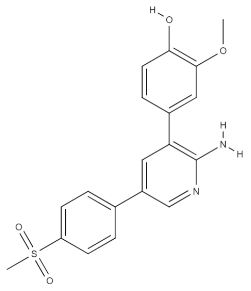

MMV010576

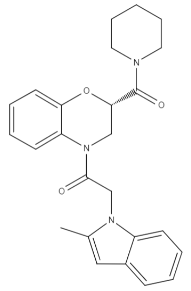

MMV020520

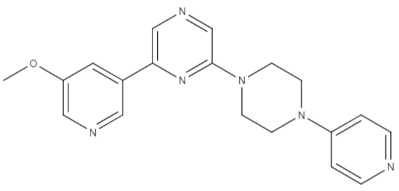

MMV085071

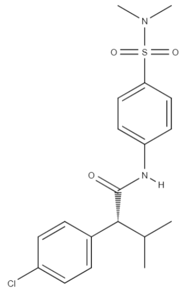

MMV020391

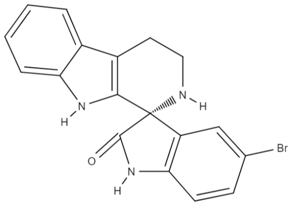

MMV006239

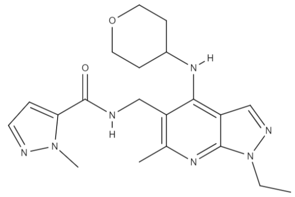

MMV026356

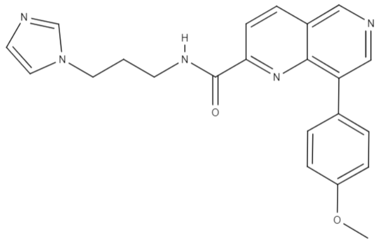

MMV020670

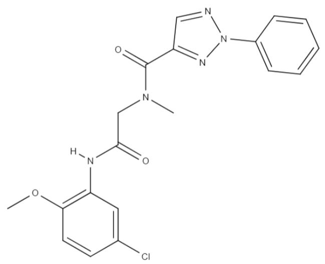

MMV020710

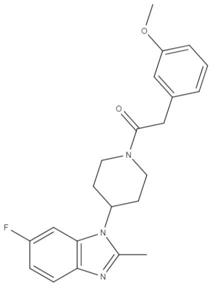

MMV020081

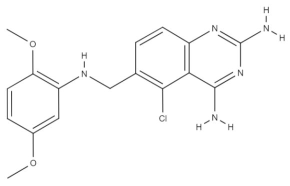

MMV675968

**Fig S1**: Structure of the 12 most effective compounds identified by *in vitro* screening using 1μM as the highest concentration, as late stage parasite inhibitors. These compound structures are obtained from PubChem (https://pubchem.ncbi.nlm.nih.gov) and MolView (http://molview.org/)

### Fig S2

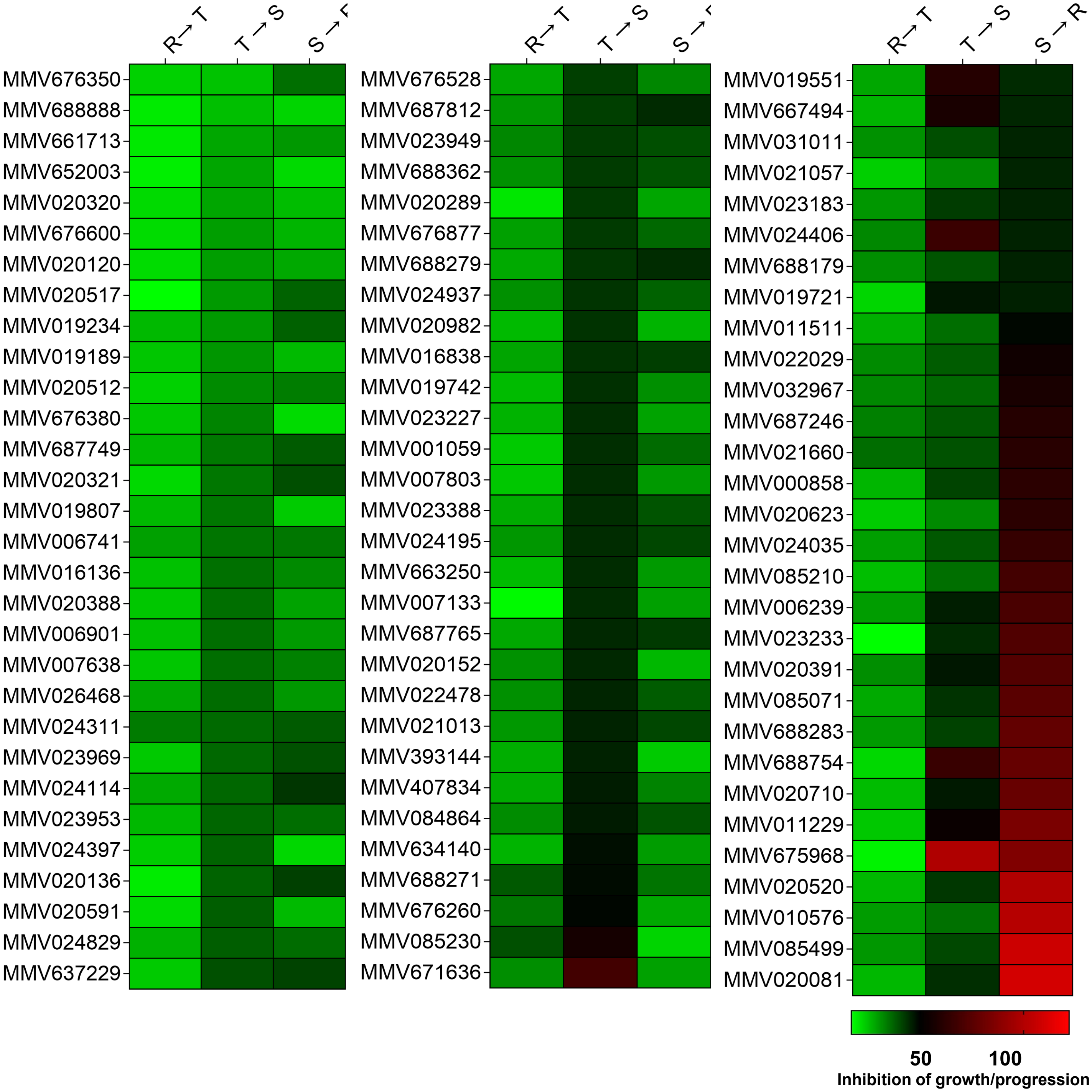


**Fig S2**: Heat Map showing parasite growth progression data of 96 molecules testing against *P. falciparum* .The data shown are mean values of two biological replicates.
