## Supplementary Figure S2 for "Whole Cell Phenotypic Screening Of MMV Pathogen Box identifies Specific Inhibitors of *Plasmodium falciparum* merozoite maturation and egress"

**Table 2: Molecules from other disease sets exhibiting antimalarial activity**

| ChEMBL ID | Pathogen Box ID | Compound Class | Disease Set | Pfal IC50 (µM) (this study) | Cytotoxicity Data (Pathogen Box dataset) | Predicted Cellular Target | Literature References |
|---|---|---|---|---|---|---|---|
| **CHEMBL227240** | MMV688514 | Benzenecarboximidamide | Kinetoplastids | 3.66 | 2.64 | No Hits | 1 |
| **CHEMBL466426** | MMV688943 | Difenoconazol | Kinetoplastids | 6.18 | 0 | No Hits | NA |
| **CHEMBL564829** | MMV676602 | Milciclib | Kinetoplastids | 6.06 | <0.26 | No Hits | 3 |
| **CHEMBL2441383** | MMV652003 | Benzamide | Kinetoplastids | 0.11 | 0.413 | Leucyl-tRNA synthetase | 34 |
| **CHEMBL1981047/ CHEMBL384575/ CHEMBL402548** | MMV676600 | Benzamide | Kinetoplastids | 0.25 | 4.83 | No Hits | 5 |
| **CHEMBL1836562** | MMV688410 | Acetamide | Kinetoplastids | 3.85 | 16.4 | No Hits | 6 |
| **CHEMBL1197423** | MMV688547 | 1H-1,2,3-Triazole | Kinetoplastids | 1.78 | >80 | DNA Targeted agent | 7,10 |
| **CHEMBL3637895** | MMV688283 | 4-Quinolinamine | Kinetoplastids | 0.50 | 8 | Potentially target Beta-hematin formation | 35 |
| **CHEMBL394241** | MMV676057 | 1-Piperazinecarboxylic acid | Kinetoplastids | 6.51 | 4.03 | No Hits | 9 |
| **CHEMBL45843** | MMV688179 | 2,2'-[2,5-Furandiylbis(3-chlor-4,1-phenylen)] diguanidin | Kinetoplastids | 0.74 | 6.75 | No Hits | 10 |
| **CHEMBL227667** | MMV688361 | Isoxazole | Kinetoplastids | 1.45 | 4.51 | No Hits | 1 |
| **CHEMBL3431105** | MMV595321 | Benzamide | Kinetoplastids | 5.59 | 0 | No Hits | 11 |
| **CHEMBL3637896** | MMV688371 | Benzamide | Kinetoplastids | 5.85 | 0 | No Hits | NA |
| **CHEMBL474899** | MMV659004 | Pyrimidine | Kinetoplastids | 2.39 | 10.2 | Methionine aminopeptidase | 13 |
| **CHEMBL3637894** | MMV688274 | 4-Morpholineacetamide | Kinetoplastids | 3.48 | 10.9 | No Hits | NA |
| **CHEMBL1197424** | MMV688407 | 1H-1,2,3-Triazole | Kinetoplastids | 2.92 | 22.9 | No Hits | 7 |

| | | | | | | | |
|---|---|---|---|---|---|---|---|
| **CHEMBL520654** | MMV688362 | 1H-Imidazole | Kinetoplastids | 0.42 | 14.3 | Binds to the DNA minor groove at AT-rich regions of DNA | 15 |
| **CHEMBL1230468** | MMV688180 | Benzenesulfonamide | Kinetoplastids | 5.31 | 0.0454 | No Hits | 2,4,8 |
| **CHEMBL1836611** | MMV688279 | 4-Quinazolinol | Kinetoplastids | 0.22 | 10 | No Hits | 6 |
| **CHEMBL238442** | MMV687706 | Piperazine | Kinetoplastids | 7.65 | 6.43 | No Hits | 9 |
| **CHEMBL413331** | MMV688271 | Guanidine | Kinetoplastids | 0.40 | 13.5 | Binds to the DNA minor groove at AT-rich regions of DNA | 10 |
| **CHEMBL3104375** | MMV689243 | 3-Pyridinamine | Kinetoplastids | 2.00 | 25.8 | Cytochrome P450 inhibitors | 16 |
| **CHEMBL1897483/ CHEMBL3183941/ CHEMBL3637899** | MMV688754 | Benzeneacetic acid | Kinetoplastids | 0.01 | 0 | No Hits | NA |
| **CHEMBL3637893** | MMV688273 | 4,6-Quinazolinediamine | Kinetoplastids | 1.61 | 9.72 | No Hits | NA |
| **CHEMBL472881** | MMV658988 | 4-Pyrimidinamine | Kinetoplastids | 2.84 | 11.3 | Methionine aminopeptidase | 13 |
| **CHEMBL3637898** | MMV688550 | Imidazo[1,2]purine | Kinetoplastids | 3.05 | 44.3 | No Hits | NA |
| **CHEMBL2097846** | MMV688889 | pyrimidin-7-amine | Tuberculosis | 8.24 | NA | No Hits | 12 |
| **CHEMBL2098256** | MMV688888 | Pyrimidine | Tuberculosis | 0.95 | NA | No Hits | 12,14 |
| **CHEMBL204432** | MMV661713 | 4-Pyrimidinamine | Tuberculosis | 0.60 | NA | Glutaminyl cyclase | 12,17 |
| **CHEMBL2109905** | MMV688936 | 2-Furancarboxamide | Tuberculosis | 6.60 | NA | No Hits | 12 |
| **CHEMBL1622353** | MMV676401 | 4-Quinazolinamine | Tuberculosis | 5.99 | 18.59 | No Hits | 18 |
| **CHEMBL2098478** | MMV676449 | 1H-Benzimidazole-1-ethanol | Tuberculosis | 8.51 | 43.2 | No Hits | 12 |
| **CHEMBL2098184** | MMV676477 | Benzamide | Tuberculosis | 0.76 | 1.34 | No Hits | 12,14 |
| **CHEMBL2098293** | MMV676512 | 1H-Imidazole-5-carboxamide | Tuberculosis | 3.66 | 3.78 | No Hits | 12 |
| **CHEMBL2109740** | MMV687703 | 1H-Benzimidazole | Tuberculosis | 4.03 | 24.98 | No Hits | 12 |

| | | | | | | | |
|---|---|---|---|---|---|---|---|
| **CHEMBL535116** | MMV023969 | Isoquinoline | Tuberculosis | 0.19 | 8.02 | No Hits | NA |
| **CHEMBL531893** | MMV021660 | Guanidine | Tuberculosis | 0.07 | 6.46 | Folate Pathway | 12,19,20 |
| **CHEMBL2098162** | MMV688122 | 2-pyrimidin-4-amine | Tuberculosis | 5.53 | 14.26 | Methionine aminopeptidase | 12,21 |
| **CHEMBL2109766** | MMV687749 | Pyrimidine | Tuberculosis | 0.85 | 29.03 | Tyrosine Kinase | 12 |
| **CHEMBL2109769** | MMV687248 | 1H-Benzimidazol-2-amine | Tuberculosis | 1.41 | 31.5 | No Hits | 12 |
| **CHEMBL2109877** | MMV688125 | Benzenesulfonamide | Tuberculosis | 4.04 | 17.69 | No Hits | 12 |
| **CHEMBL561057** | MMV687273 | SQ-109 , In clinical trials 2012 TB | Tuberculosis | 3.29 | 7.1 | HMG-CoA reductase pathway? | 22,23,20,24, 25 |
| **CHEMBL580032** | MMV024311 | 1H-Indole | Tuberculosis | 0.45 | 12.71 | No Hits | 12 |
| **CHEMBL2313136** | MMV687807 | Benzamide | Tuberculosis | 2.47 | 0.65 | No Hits | 12,26,27 |
| **CHEMBL3637864** | MMV687812 | 2-Pyrazinecarboxamide | Tuberculosis | 0.26 | 3.86 | No Hits | 28 |
| **CHEMBL1397830** | MMV676411 | Propanamide | Tuberculosis | 5.02 | 29.29 | No Hits | 12 |
| **CHEMBL530275** | MMV021013 | 4-pyrimidinamine | Tuberculosis | 0.72 | NA | Methionine aminopeptidase | 12 |
| **CHEMBL2109928** | MMV687765 | Pyrimidine | Tuberculosis | 0.47 | 14.08 | Tyrosine Kinase | 12 |
| **CHEMBL3637867** | MMV495543 | Benzamide | Tuberculosis | 0.26 | 47.64 | No Hits | NA |
| **CHEMBL1242290** | MMV675993 | | Cryptosporidiosis | 3.93 | 1.63 | No Hits | 29 |
| **CHEMBL88430** | MMV675968 | | Cryptosporidiosis | 0.07 | 3.44 | Dihydrofolate reductase | 30,31,32 |
| **CHEMBL1626/315157/1620822** | MMV637229 | 2-pyrimidin-4-amine | Hookworm Trichuriasis | 0.01 | 9.28 | No Hits | NA |
| **CHEMBL3637906** | MMV668727 | Terpyridin-2'-amine | L. Filariasis-Onchocerciasis | 3.64 | >80 | No Hits | NA |
| **CHEMBL3637912** | MMV676063 | oxy-phenyl urea | L. Filariasis-Onchocerciasis | 3.16 | 5.77 | No Hits | NA |

| | | | | | | | |
|---|---|---|---|---|---|---|---|
| **CHEMBL3637903** | MMV671636 | quinolinone | L. Filariasis-Onchocerciasis | 0.01 | >80 | Mitochondria cytochrome $bc_1$ complex | NA |
| **CHEMBL3637922** | MMV688761 | Benzamide | Schistosomiasis | 3.46 | 6.94 | No Hits | NA |
| **CHEMBL3637921** | MMV688763 | pyridazinone | Schistosomiasis | 2.17 | 1.55 | No Hits | NA |
| **CHEMBL3637924** | MMV688766 | 5-methoxyphenol | Schistosomiasis | 1.71 | >80 | No Hits | 36 |
| **CHEMBL3637923** | MMV688552 | 1H-Pyrazole-1-carboxamide | Schistosomiasis | 4.49 | 30.7 | No Hits | NA |
| **CHEMBL3637929** | MMV688509 | 1-Piperidinecarboxylic acid | Toxoplasmosis | 5.55 | >50 | No Hits | NA |
| **CHEMBL190228** | MMV688703 | Pyridine | Toxoplasmosis | 1.39 | >50 | cGMP-dependent protein kinase | 33 |
